# Schizophrenia patients show aberrant brain dynamics associated with gestalt-perception and corollary discharge: Reflections from ERP and fMRI findings

**DOI:** 10.1101/837831

**Authors:** Arun Sasidharan, Ajay Kumar Nair, Vrinda Marigowda, Ammu Lukose, John P John, Bindu M Kutty

## Abstract

**Background:** Schizophrenia is a disorder of higher mental attributes, and is characterized by psychotic symptoms that are believed to involve a basic inability to make valid predictions about expected sensations and experiences. These have been reported separately while monitoring either externally generated environmental patterns (e.g. gestalt-perception) or self-generated sensory experiences (e.g. corollary-discharge). As the pathophysiology behind predictive dysfunction is better viewed as an aberration in brain’s functional synchrony, a whole brain assessment using electroencephalographic (EEG) event related potential (ERP) and functional magnetic resonance imaging (fMRI) techniques, would offer a wider perspective to brain network abnormalities in schizophrenia.

**Method:** We used our lab-developed game-based task which presents degraded two-tone images to assess gestalt-perception, and simultaneously alters the congruency between participant’s button-press response and its auditory feedback to assess corollary-discharge. In both patients with schizophrenia and age-matched healthy controls, we explored event-related changes in an EEG-ERP study (n=21 each) and whole brain functional connectivity changes in a fMRI study (patients,n=12; controls,n=16), using the same task.

**Results:** Patients showed reduced event-related EEG dynamics during both the error-prediction conditions (gestalt-perception and corollary-discharge), which include reduction in average waveforms (around N170 and N1-P2 complex, respectively) and altered theta dynamics (power and phase). Source-level EEG measures were clustered around the cingulo-insular network. fMRI functional connectivity analysis also found the abnormality in these brain regions, forming significantly weak connections with right insular/opercular cortex.

**Conclusions:** This is the first study to explore ‘thalamo-cortical dysfunction hypothesis’ in schizophrenia by integrating prediction-error-coding during perception (‘gestalt-perception’) and action (‘corollary-discharge’) using two neuroimaging modalities (EEG-ERP and fMRI). Besides adding to the knowledgebase of schizophrenia research, our novel task design and findings on theta-oscillation could benefit in the development of effective neuromodulatory therapeutic tools for patients with schizophrenia such as neurofeedback and transcranial brain stimulation.

## Introduction

Recent advances in neuroscience, have identified schizophrenia as a pan-cerebral illness, affecting different brain-regions, at varying brain-states, and resulting in diverse brain-function abnormalities. Many attempts have failed to identify dysfunctional brain mechanisms that explain the varied symptoms presented by patients with schizophrenia (PSZ). This prompted National Institute of Mental Health (NIMH) to come up with Research Domain Criteria (RDoC)^1^ - a research classification system for mental disorders based on dimensions of neurobiology and observable behaviour. It advocates integration of neurobiology-based conceptual framework and clinically practical assessments designed to provide reliable biomarkers of cognitive dysfunction. One such framework suggests failure in brain’s prediction mechanism(s) among PSZ^2^. This may be attributable to faulty functional interaction between brain areas, thus preventing appropriate interpretation of ‘noisy’ information bombarded from outside world. Thereby, PSZ misinterpret what they see, hear or experience in their environment, and this interferes with their ability to think clearly and manage emotions.

One such ‘de-noising’ mechanism is top-down prediction-error-coding, where higher brain centers (association areas) predict incoming information, which is then compared with bottom-up stimulus-bound signals from lower centers (sensory areas)^3^. Discrepancy between bottom-up sensory stimulus and top-down predictive signals is coded as ‘prediction-errors’, and these generally form the bulk of brain signals during wake-state cognition. Such prediction-error-coding requires multiple levels of abstraction, which is known to involve medial prefrontal cortex and temporo-parietal junction, besides many other brain regions^4^. Studies on schizophrenia have confirmed failure in prediction-error-coding^2, 5–7^. Such studies examined prediction-error-coding while observing externally generated sensory patterns (gestalt-perception)^8, 9^ or while experiencing self-generated stimuli (corollary-discharge)^10, 11^. Gestalt-perception involves grouping of local stimulus features from lower centers based on matching with predictions of higher centers, to form an abstract percept (e.g. scenery). Corollary-discharge involves communication between higher motor areas and sensory areas to predict sensory outcome of self-generated action (e.g. speech). In both cases, sensory attenuation occurs if prediction-error is minimal, i.e., when perceiving an ‘abstract sense’ (gestalt-perception) or a sense of ‘self-generation’ (corollary-discharge). In schizophrenia, aberrant prediction-errors are generated that lead to inappropriate salience and thereby misinterpretations^2^. As cortico-thalamo-cortical loop is essential for the dynamic formation of inter-cortical networks implicated in prediction-error-coding, this circuitry could be altered in schizophrenia and forms a putative target for research^2, 12^.

Evidence for ‘thalamo-cortical dysfunction hypothesis’ of schizophrenia^12–15^, is rooted from several isolated research studies in schizophrenia. Firstly, whole-night sleep studies have reported sleep spindles deficits with or without sleep architecture abnormalities, suggesting aberrant thalamo-cortical interaction^16, 17^. Secondly, deficits in event related potentials (ERPs) has been interpreted as deficits in thalamus mediated filtering of novel/salient stimuli^18, 19^. Thirdly, neuroimaging studies have reported thalamic abnormality either in terms of morphology^20^ or activation pattern during various tasks^21^.

However, as this circuitry handles prediction-error-coding during perception and action differently^3^, there is a need to integrate both these aspects for exploring ‘thalamo-cortical dysfunction hypothesis’ in schizophrenia. Accordingly, the current study integrated two tasks (‘gestalt-perception’ - perceiving whole images from incomplete visual patterns; and ‘corollary-discharge’ - distinguishing ‘self-generated’ from ‘other-generated’ stimuli) and used two neuroimaging modalities (EEG-ERP and functional magnetic resonance imaging (fMRI)). The study hypothesized that PSZ would show varying degrees of abnormalities in task-related EEG-ERP and task-related fMRI connectivity as markers of altered ‘predictive error’, and thence the dysfunctional thalamo-cortical network.

## Methods and Materials

The study was carried out with approval from Institute Ethics Committee, thus conforming to the ethical standards laid down in the 1964 Declaration of Helsinki. Written informed consent was obtained from all participants (and their legally qualified representatives in the case of PSZ) prior to enrolling them into the study.

**Note:** Please refer to supplementary methodology for more details.

### Participants

Healthy control subjects (HCS) were recruited by word of mouth, and PSZ were recruited from the out-patient department by purposive sampling. Participants were matched for age and gender (Table-1).

**Table 1:** Sociodemographic details of participants of ERP and fMRI study.

|  |  | Healthy Control Subjects | Patients with Schizophrenia | Statistics |
| --- | --- | --- | --- | --- |
| **Sex** | ERP | M=12; F=9; T=21 | M=14; F=5; T=19 | X-squared=1.200;p=0.273 |
|  | fMRI | M=11; F=5; T=16 | M=10; F=2; T=12 | X-squared=0.778;p=0.378 |
| **Age (yrs)** | ERP | 29.14±4.79 (20-42) | 27.84±7.14 (19-44) | t=0.647;p=0.521;CI=-2.7 to 5.0 |
|  | fMRI | 24.56±4.68 (18-32) | 27.25±4.41 (19-33) | t=-1.554;p=0.142;CI=-0.5 to 5.9 |
| **Education (yrs)#** | ERP | 19.05±2.23 (13-24) | 11.94±3.37 (3-17) | t=7.374;p<0.001;CI=5.3 to 8.9 |
|  | fMRI | 14.70±1.57 (13-18) | 14.36±3.38 (9-18) | t=0.297;p=0.712;CI=-2.5 to 1.7 |
| **Age of Illness onset (yrs)** | ERP |  | 25.09±7.03 (17-43.7) |  |
|  | fMRI |  | 24.42±3.45 (17-29) |  |
| **Illness duration (months)** | ERP |  | 34.86±30.17 (3-120) |  |
|  | fMRI |  | 34.67±25.01 (8-84) |  |
| **Antipsychotic exposure (months)** | ERP |  | 15.08±20.89 (0-72) |  |
|  | fMRI# |  | 22.10±24.98 (0-72) |  |
| **Treatment status** | ERP |  | uPSZ=7; mPSZ=12 |  |
|  | fMRI |  | uPSZ=2; mPSZ=10 |  |
| **NOTE:** All data shown as: Mean ± SD (min-max). t statistic and p-value for group comparisons obtained by non-parametric permutation based t-test using 10000 iterations. CI: 95% confidence interval of the mean difference obtained using bootstrapping statistics on 10000 re-samples. Grey shading represents statistically significant between groups at p<0.05. # Data not available for all participants. uPSZ – unmedicated patients with schizophrenia; mPSZ - medicated patients with schizophrenia. |  |  |  |  |

### Task design

‘Prediction-error-coding’ was assessed using our custom-developed task (‘ANGEL’ or Assessing Neurocognition via Gamified Experimental Logic^22^) that helps in simultaneous assessment of multiple ERPs in a single session. A shortened version was used for fMRI study.

### EEG recording

All recordings were done in a sound attenuated chamber, using a 70-channel Neuroscan SynAmps2 acquisition system (Compumedics, Charlotte, USA), [24bits resolution; 1000Hz sampling rate; 0.1-100Hz high-pass filter; no notch-filters] and gel-based sintered-silver electrode caps (Neuroscan-Quik caps).

### ERP analysis

Offline analysis was performed using EEGLAB v13 toolbox^23^ in MATLAB 2013a. Automated artefact removal was employed using artefact subspace reconstruction (ASR) algorithm^24^ implemented as a plugin in EEGLAB. ERPs were obtained by averaging of 3000ms pre-processed EEG epochs starting 1000ms before the onset of gestalt images (gestalt-perception) or polysyllabic feedback sound (corollary-discharge). A low pass finite impulse response (FIR) filter of 35Hz was applied to ERP data. Electrode for initial exploration of ERP as well as time-frequency analysis (ERSP and ITC) was chosen based on those reported in previous studies. Accordingly, PO7/PO8 electrodes were used to observe the prominent N170-P200 complex during gestalt-perception, and FCz/Cz electrodes were used to observe the N100-P200 suppression during corollary-discharge. However, time points to extract subject level differences were decided based on the group-level analysis at these electrodes.

### Event-Related Spectral Perturbations (ERSP) and Inter-Trial Coherence (ITC) analysis

ERSP (strength of power modulations across trials) and ITC (measure of strength of phase alignment across trials) were computed using wavelet-transform routines of EEGLAB. Due to computational limitations, we restricted to a frequency window of 3-30Hz for all channels.

### EEG Source level analysis

To further examine the brain sources of ERP signals, first an independent component analysis (ICA) was performed, and then the location of each independent component (IC) was computed using DIPFIT plugin of EEGLAB. ERP, ERSP and ITC computed from ICs of all subjects, were each subjected to a probabilistic multi-subject inference using Measure Projection Analysis (MPA) implemented in MPT (Measure Projection Toolbox) plugin of EEGLAB^25^.

### fMRI acquisition

fMRI was done using 3-Tesla wide-bore MRI scanner (Skyra, Seimens, Germany) with 20-channel head-coil. A total of 445 scans were obtained (after discarding first 2 dummy scans).

### fMRI Analysis

fMRI images were pre-processed using Statistical Parametric Mapping (SPM v12) toolbox routines and further functional connectivity analysis using CONN v14p toolbox^26^, running on MATLAB 2013a. We used 136 default regions of interest (ROIs; 95 cortical and 51 subcortical) and used bivariate correlations as connectivity estimations during error prediction trials (both gestalt-perception and corollary-discharge tasks combined).

### Statistical analysis

All statistical tests were implemented using statistical functions that were part of the EEGLAB v13 toolbox (for EEG data) or SPM12 and CONNv14p (for fMRI data), implemented in MATLAB 2013a software. Permutation-based two-way mixed design analysis of variance (ANOVA) was used to test significant differences in EEG data, after multiple comparison correction using false discovery rate (FDR). Statistical comparison of demographic parameters was computed by non-parametric permutation based t-test using 10000 iterations, and 95% confidence interval of mean difference was obtained using bootstrap statistics on 10000 re-samples. Statistical significance between groups was set at p<0.05.

## Results

### Scalp EEG dynamics of Prediction-error-coding

#### a) Gestalt-perception

PSZ showed reduced N170 (175-200ms), and significant scalp topography deficits over frontal and parieto-occipital sites, during all image types (Fig-1). PSZ also showed significant difference in P200 and later ERP components (250-550ms) for ‘Shape absent’ images, and significant scalp topography deficits over fronto-central sites mostly for ‘Shape absent’ and ‘Shape present’ images. Both these time points also showed most significant between-condition changes in both groups.

**Fig-1:**
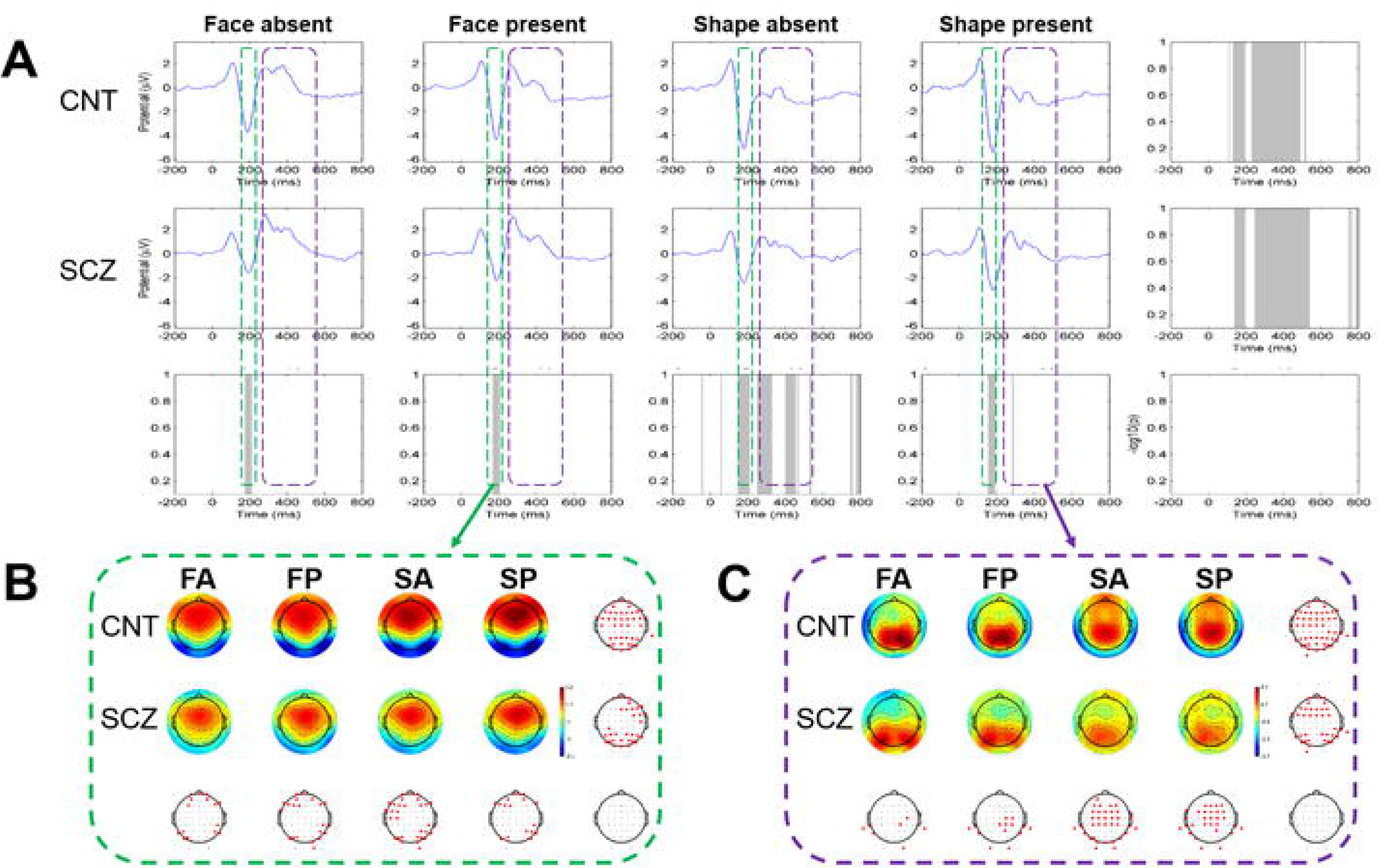
Event related potential (ERP) during gestalt-perception in healthy controls (CNT) and patients with schizophrenia (SCZ). (A) Grand average ERPs from 21 subjects of either groups for ‘Face absent’ (FA), ‘Face present’ (FP), ‘Shape absent’ (SA) and ‘Shape present’ (SP) conditions, at PO8 electrode site. Statistical results are shown as grey shaded areas in the lower and right panels. Scalp topography of the mean ERPs over 2 time points that showed between-group statistical significance: (B) 175-200ms, and (C) 250-550ms. Statistically significant electrode sites are shown as dark red spots on unfilled scalp maps. Statistical analysis used permutation based (800 permutations) two-way mixed design ANOVA and post-hoc t-tests with FDR correction at <0.05.

ERSP changes included an increase in theta (3-8Hz) power followed by reduction in beta (13-30Hz) power at PO8 electrode, in both groups (Fig-2). PSZ showed significant deficit in theta power at the post-image period (0-300ms) mostly for ‘Shape absent’ and ‘Shape present’ images, and significant scalp topographic deficits over frontal sites. Additionally, theta power reduction occurring in occipital sites were also significantly less in PSZ. ITC analysis also showed significant deficit for PSZ in theta phase resetting for ‘Shape absent’ images at the peri-stimulus time (−100 to 200ms), and scalp topographic deficits mainly over frontal sites for all image types (Fig-3).

**Fig-2:**
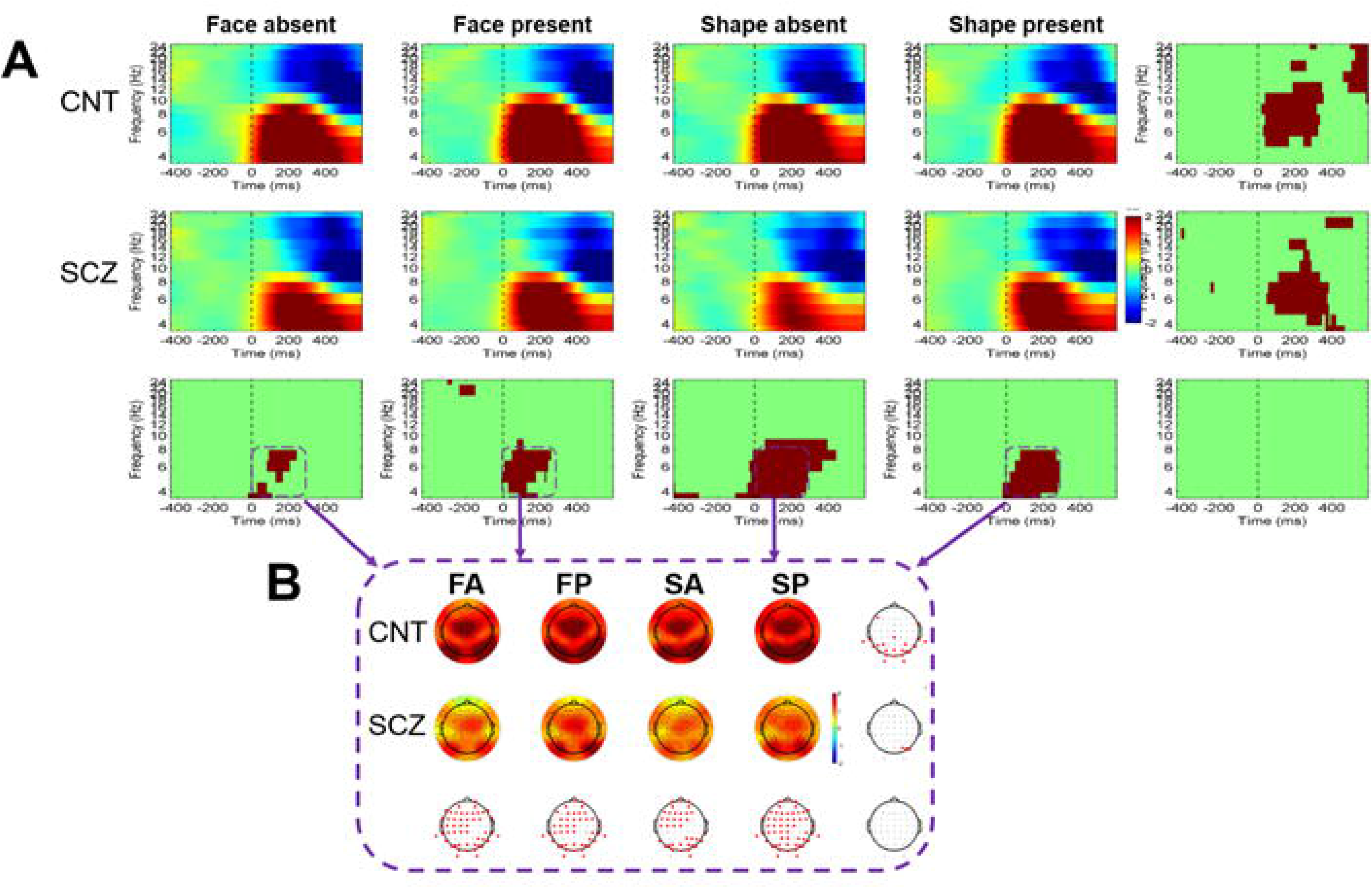
Event related spectral perturbation (ERSP) during gestalt-perception in healthy controls (CNT) and patients with schizophrenia (SCZ). (A) Grand average ERSPs from 21 subjects of either groups for ‘Face absent’ (FA), ‘Face present’ (FP), ‘Shape absent’ (SA) and ‘Shape present’ (SP) conditions, at PO8 electrode site. Statistical results are shown as grey shaded areas in the lower and right panels. (B) Scalp topography of the mean ERSPs in the theta band (3-8Hz) over the time point that showed between-group statistical significance (0-300ms). Statistically significant electrode sites are shown as dark red spots on unfilled scalp maps. Statistical analysis used permutation based (800 permutations) two-way mixed design ANOVA and post-hoc t-tests with FDR correction at <0.05.

**Fig-3:**
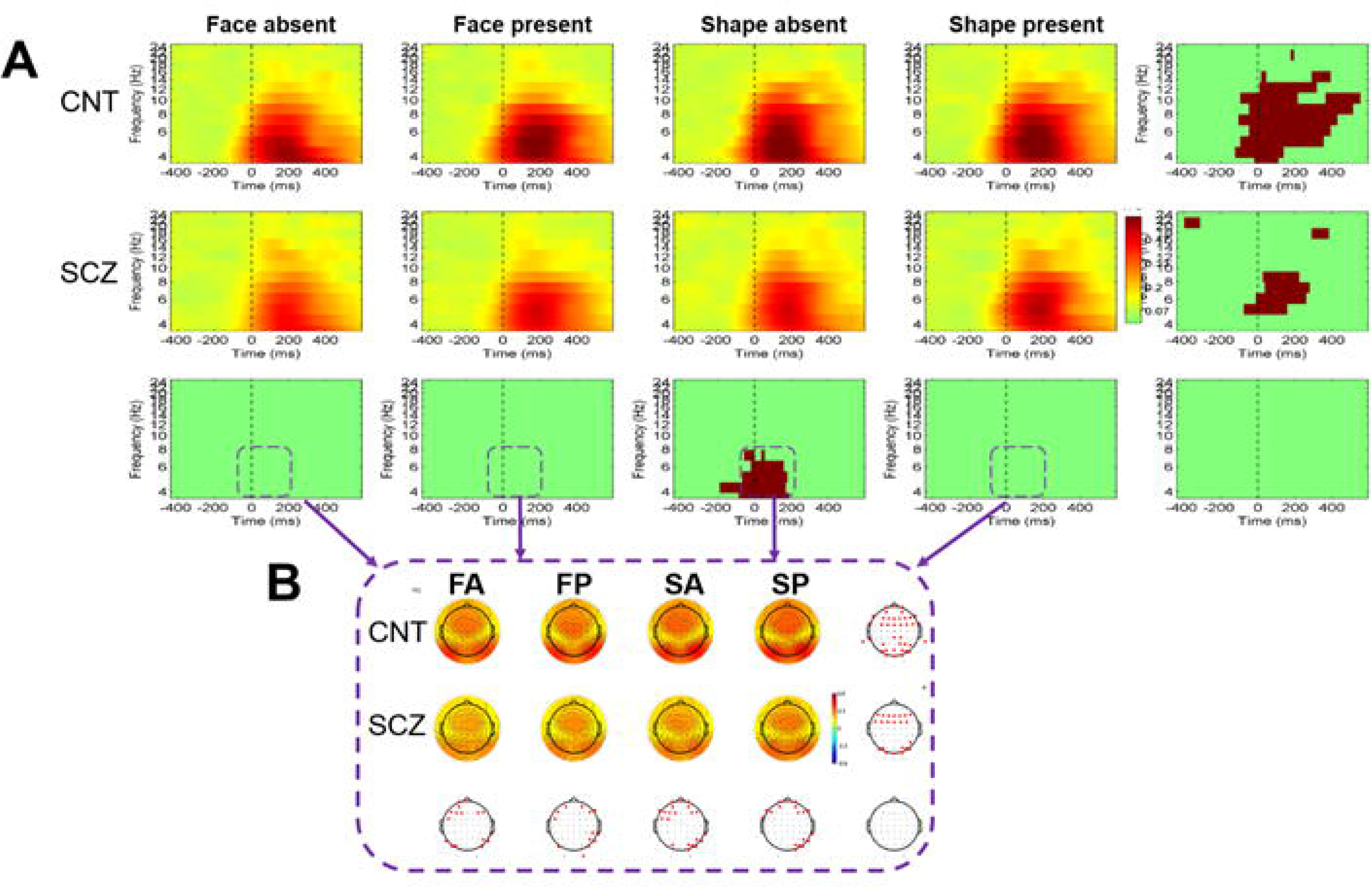
Inter-trial coherence (ITC) during gestalt-perception in healthy controls (CNT) and patients with schizophrenia (SCZ). (A) Grand average ITCs from 21 subjects of either groups for ‘Face absent’ (FA), ‘Face present’ (FP), ‘Shape absent’ (SA) and ‘Shape present’ (SP) conditions, at PO8 electrode site. Statistical results are shown as grey shaded areas in the lower and right panels. (B) Scalp topography of the mean ITCs in the theta band (3-8Hz) over the time point that showed between-group statistical significance (−100 to 200ms). Statistically significant electrode sites are shown as dark red spots on unfilled scalp maps. Statistical analysis used permutation based (800 permutations) two-way mixed design ANOVA and post-hoc t-tests with FDR correction at <0.05.

#### b) Corollary-discharge

PSZ showed reduced ERP for ‘CDon’ condition at three time-periods (−200 to −150ms, −100 to −50ms and 50 to 100ms) and no difference for ‘CDoff’ condition, compared to HCS (Fig-4). Scalp topographic deficits were mainly over central electrodes.

**Fig-4:**
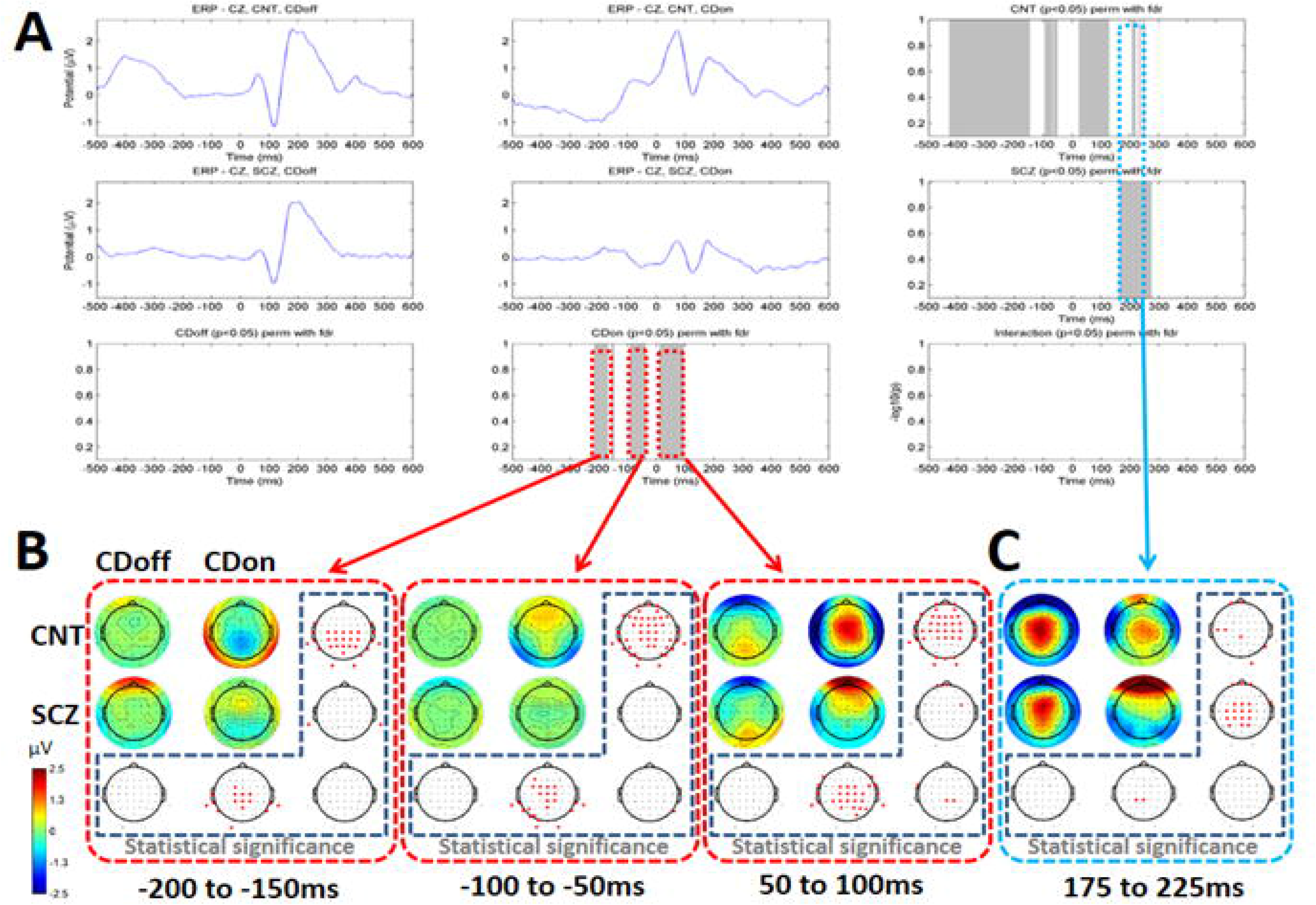
Event related potential (ERP) during corollary discharge mechanism in healthy controls (CNT) and patients with schizophrenia (SCZ). (A) Grand average ERPs from 21 subjects of either groups for ‘CDon’ and ‘CDoff’ conditions, at Cz electrode site. Statistical results are shown as grey shaded areas in the lower and right panels. Scalp topography of the mean ERPs: (B) over 3 time points that showed between-group statistical significance, and (C) over one time point that showed between-condition statistical significance. Statistically significant electrode sites are shown as dark red spots on unfilled scalp maps. CDon/CDoff-corollary discharge present or absent respectively. Statistical analysis used permutation based (800 permutations) two-way mixed design ANOVA and post-hoc t-tests with FDR correction at <0.05.

ERSP changes were most prominent around theta frequency and showed significant main effects (both between-group and between-condition) as well as interaction effects at two time-periods (−440ms to −280ms and −160 to 60ms), with scalp topography deficits at most electrode sites (Fig-5). The difference in theta-ERSP between ‘CDoff’ and ‘CDon’ conditions was also significantly lower among PSZ, at both the time-periods (Table-2).

**Fig-5:**
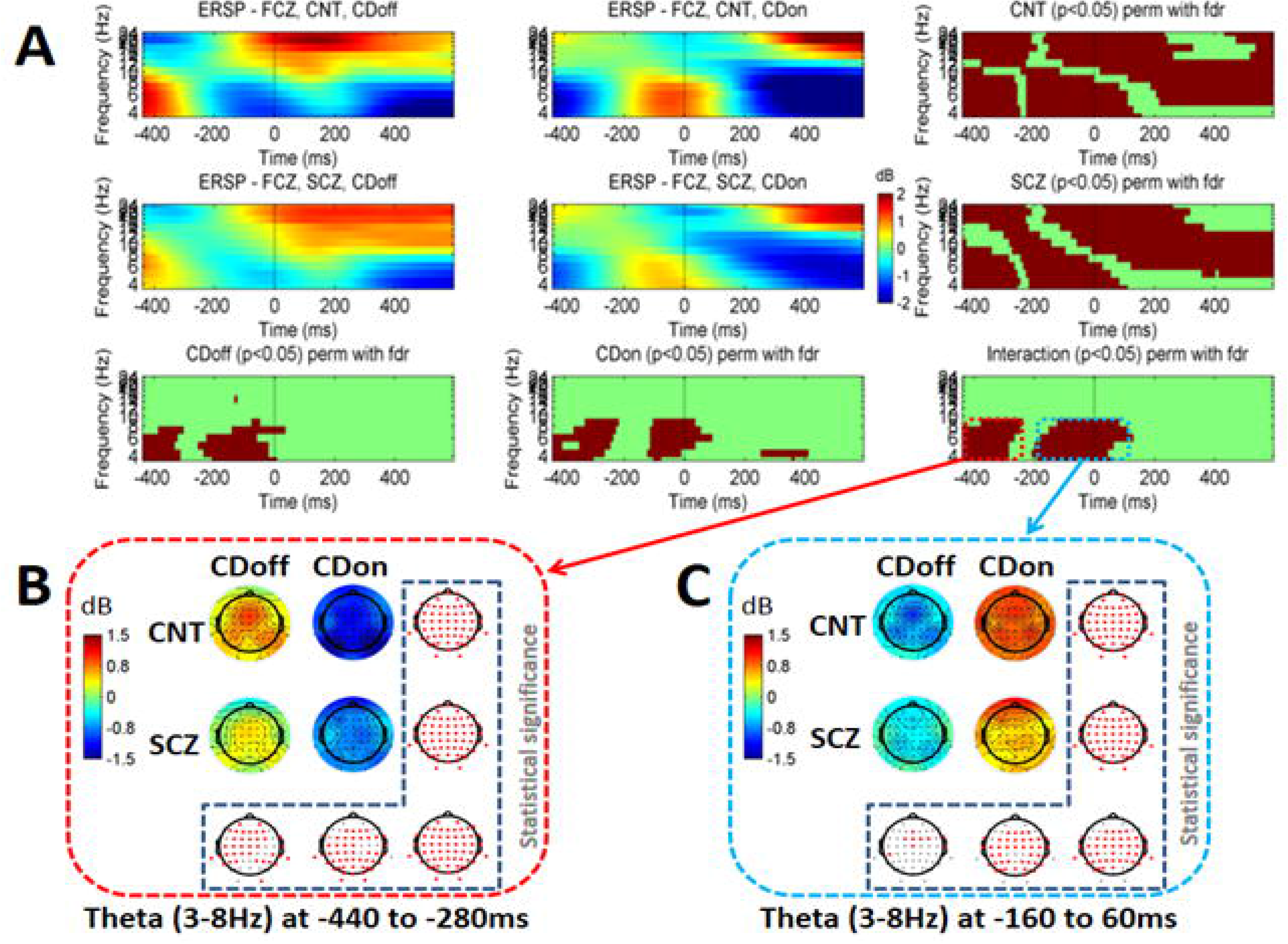
Event related spectral perturbation (ERSP) during corollary discharge mechanism in healthy controls (CNT) and patients with schizophrenia (SCZ). (A) Grand average ERSPs from 21 subjects of either groups for ‘CDon’ and ‘CDoff’ conditions, at FCz electrode site. Statistical results are shown as dark-red shaded areas in the lower and right panels. (B & C) Scalp topography of the mean ERSPs in the theta band (3-8Hz) over 2 time points that showed statistically significant group-condition interaction effect. Statistically significant electrode sites are shown as dark red spots on unfilled scalp maps. CDon/CDoff-corollary discharge present or absent respectively. Statistical analysis used permutation based (800 permutations) two-way mixed design ANOVA and post-hoc t-tests with FDR correction at <0.05.

**Table 2:** Between condition event related spectral perturbation (ERSP) differences during corollary.

| Time points evaluated | CNT | SCZ | Statistics |
| --- | --- | --- | --- |
| -440ms to -280ms* | 2.212 ± 0.519 | 1.283 ± 0.637 | t = 4.96; p < 0.001;<br>CI = 0.584 to 1.259 |
| -160 to 60ms* | -1.979 ± 0.718 | -0.973 ± 0.643 | t = -4.052; p < 0.001;<br>CI = -0.607 to -1.392 |
| **NOTE:** All data shown as: Mean ± SD. t statistic and p-value obtained by non-parametric permutation based t-test using 10000 iterations. CI: 95% confidence interval of the mean difference obtained using bootstrapping statistics on 10000 re-samples. *Statistically significant between groups at p<0.05. |  |  |  |

discharge task.

PSZ showing decreased theta-ITC values during ‘CDon’ condition at two time-periods (−200 to 400ms and 400 to 600ms), which was also distributed throughout scalp (Fig-6).

**Fig-6:**
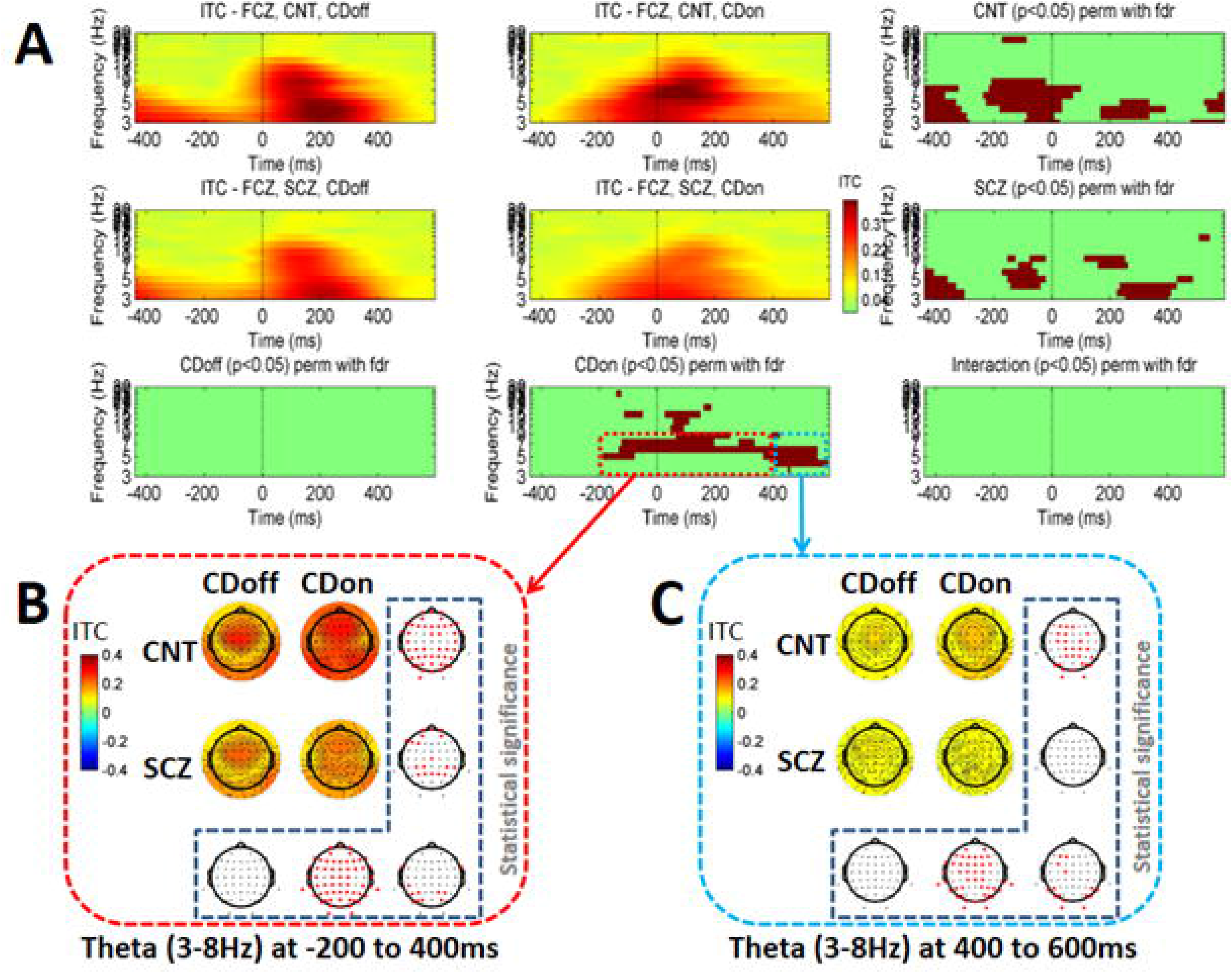
Inter-trial coherence (ITC) during corollary discharge mechanism in healthy controls (CNT) and patients with schizophrenia (SCZ). (A) Grand average ITCs from 21 subjects of either groups for ‘CDon’ and ‘CDoff’ conditions, at FCz electrode site. Statistical results are shown as dark-red shaded areas in the lower and right panels. (B & C) Scalp topography of the mean ITCs in the theta band (3-8Hz) over 2 time points that showed statistically significant group-condition interaction effect. Statistically significant electrode sites are shown as dark red spots on unfilled scalp maps. CDon/CDoff-corollary discharge present or absent respectively. Statistical analysis used permutation based (800 permutations) two-way mixed design ANOVA and post-hoc t-tests with FDR correction at <0.05.

See Table-3 for task performance.

**Table 3:** Performance measures during the ‘ANGEL’ task.

|  | Healthy Controls<br>(CNT) | Schizophrenia Patients<br>(SCZ) | Statistics |
| --- | --- | --- | --- |
| **ANGEL game level 2** |  |  |  |
| **Accuracy (%)**<br>(CNT,n=21;SCZ,n=21) | 89.39 ± 8.26<br>(74.3-100.5) | 88.30 ± 13.16<br>(36.5-99.3) | t statistic = -0.315;<br>p-value = 0.799;<br>CI = -8.1 to 4.8 |
| **Speed\* (in ms)**<br>(CNT,n=21;SCZ,n=21) | 302.11 ± 19.74<br>(270.1-332.2) | 363.26 ± 44.49<br>(299.8-428.2) | t statistic = 4.924;<br>**p-value = 0.000;**<br>CI = 41.0 to 81.2 |
| **ANGEL game level 3** |  |  |  |
| **Accuracy\* (%)**<br>(CNT,n=21;SCZ,n=21) | 72.24 ± 4.71<br>(59.3-77.5) | 55.93 ± 17.56<br>(20.0-79.3) | t statistic = -4.480;<br>**p-value = 0.000;**<br>CI = -23.9 to -8.9 |
| **Speed (in ms)**<br>(CNT,n=21;SCZ,n=21) | 390.99 ± 39.25<br>(302.4-452.6) | 411.99 ± 46.43<br>(340.9-501.4) | t statistic = 1.617;<br>p-value = 0.129;<br>CI = -4.0 to 47.1 |
| **NOTE:** Measures evaluated from 400 trials per subject, for each game level. All data shown as: Mean ± SD (min-max). t statistic and p-value obtained by non-parametric permutation based t-test using 10000 iterations. CI: 95% confidence interval of the mean difference obtained using bootstrapping statistics on 10000 re-samples. *Statistically significant between groups at p<0.05. |  |  |  |

### EEG Source dynamics of Prediction-error-coding

For convenience of reporting, clusters are named here based on the largest contributing brain region within each cluster.

#### a) Gestalt-perception (Fig-7)

**Fig-7:**
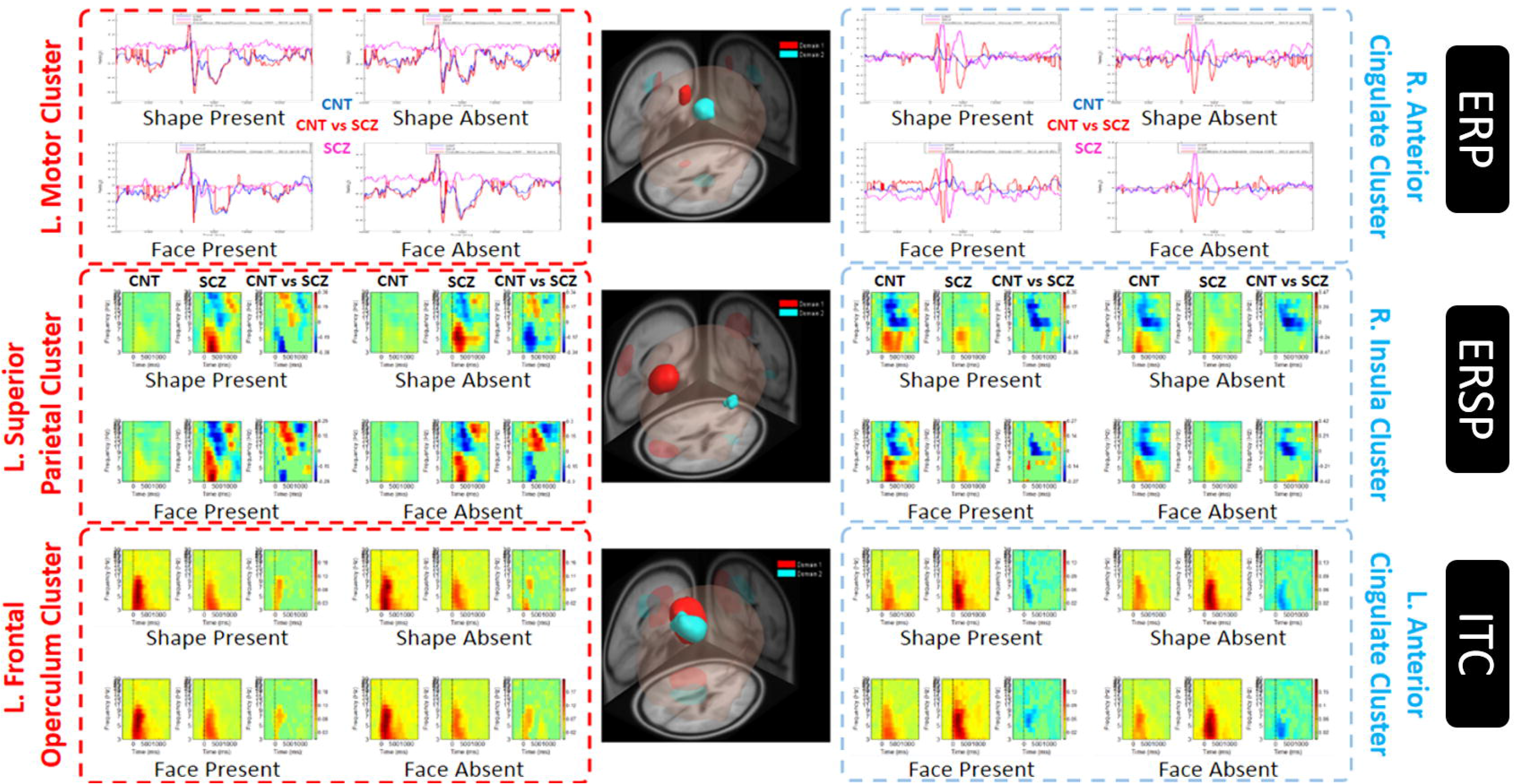
**EEG source clusters identified during gestalt-perception task using measure projection analysis** based on: (A) ERP, (B) ERSP and (C) ITC measures. Two clusters were identified for each measure, first shown on the right side (red) while second on the left side (light blue). CNT-controls; SCZ-schizophrenia. Permutation-based statistics with FDR correction set at p<0.05.

Using ERP measures, 2 dipole-clusters were localised, a left motor cluster and a right anterior cingulate cluster. The former cluster produced significantly larger ERP changes in HCS, while the latter cluster produced larger ERP changes among PSZ. Using ERSP measures, 2 dipole-clusters were localised, a left superior parietal cluster and a right insular cluster. PSZ showed significant theta power increase and beta power decrease in the former cluster, whereas showed lower dynamics in the latter cluster. Using ITC measures, 2 other clusters were identified, a left frontal opercular cluster and a left anterior cingulate cluster. Both clusters showed significant changes in theta coherence, with PSZ showing lower values in the former cluster whereas higher values in the latter cluster.

#### b) Corollary-discharge (Fig-8)

**Fig-8:**
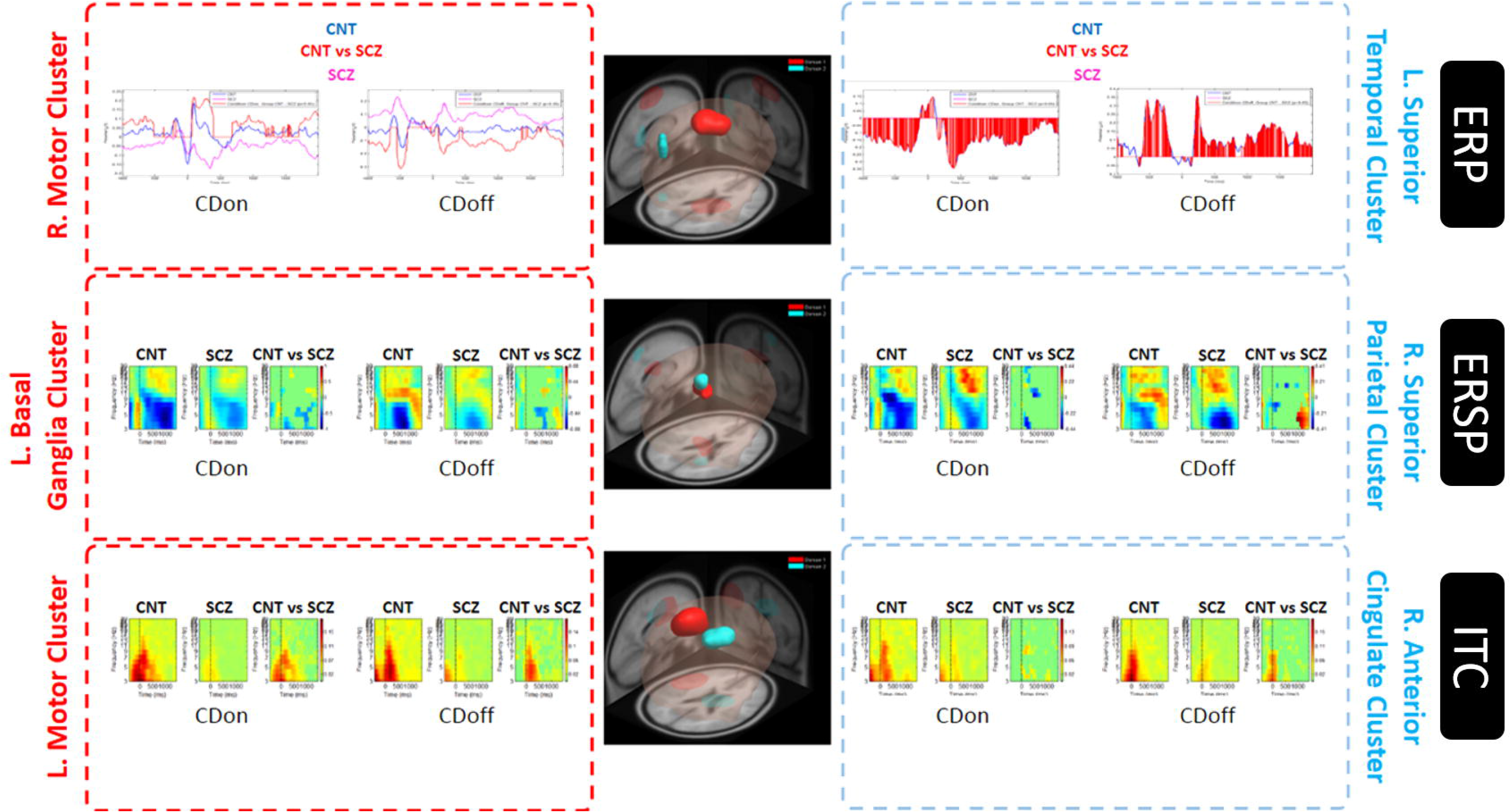
**EEG source clusters identified during corollary discharge task using measure projection analysis** based on: (A) ERP, (B) ERSP and (C) ITC measures. Two clusters were identified for each measure, first shown on the right side (red) while second on the left side (light blue). CNT-controls; SCZ-schizophrenia. Permutation-based statistics with false discovery rate correction set at p<0.05. Corollary discharge present (CDon) and absent (CDoff).

Using ERP measures, 2 dipole-clusters were localised, a right motor cluster and a left superior temporal cluster. Both groups showed significantly different ERPs generated in the former cluster, whereas PSZ showed significantly lower ERP in the latter cluster. Using ERSP measures, 2 dipole-clusters were localised, a left basal ganglia cluster and a right superior parietal cluster. Both clusters showed significantly reduced theta power dynamics in ‘CDon’ as well as ‘CDoff’ trials among PSZ. Using ITC measures, 2 other dipole-clusters were localised, a left motor cluster and a right cingulate cluster. Among PSZ, both clusters showed significantly reduced peri-stimulus theta phase synchrony.

### fMRI connectivity of Prediction-error-coding

#### a) ROI-ROI analysis (Fig-9)

**Fig-9:**
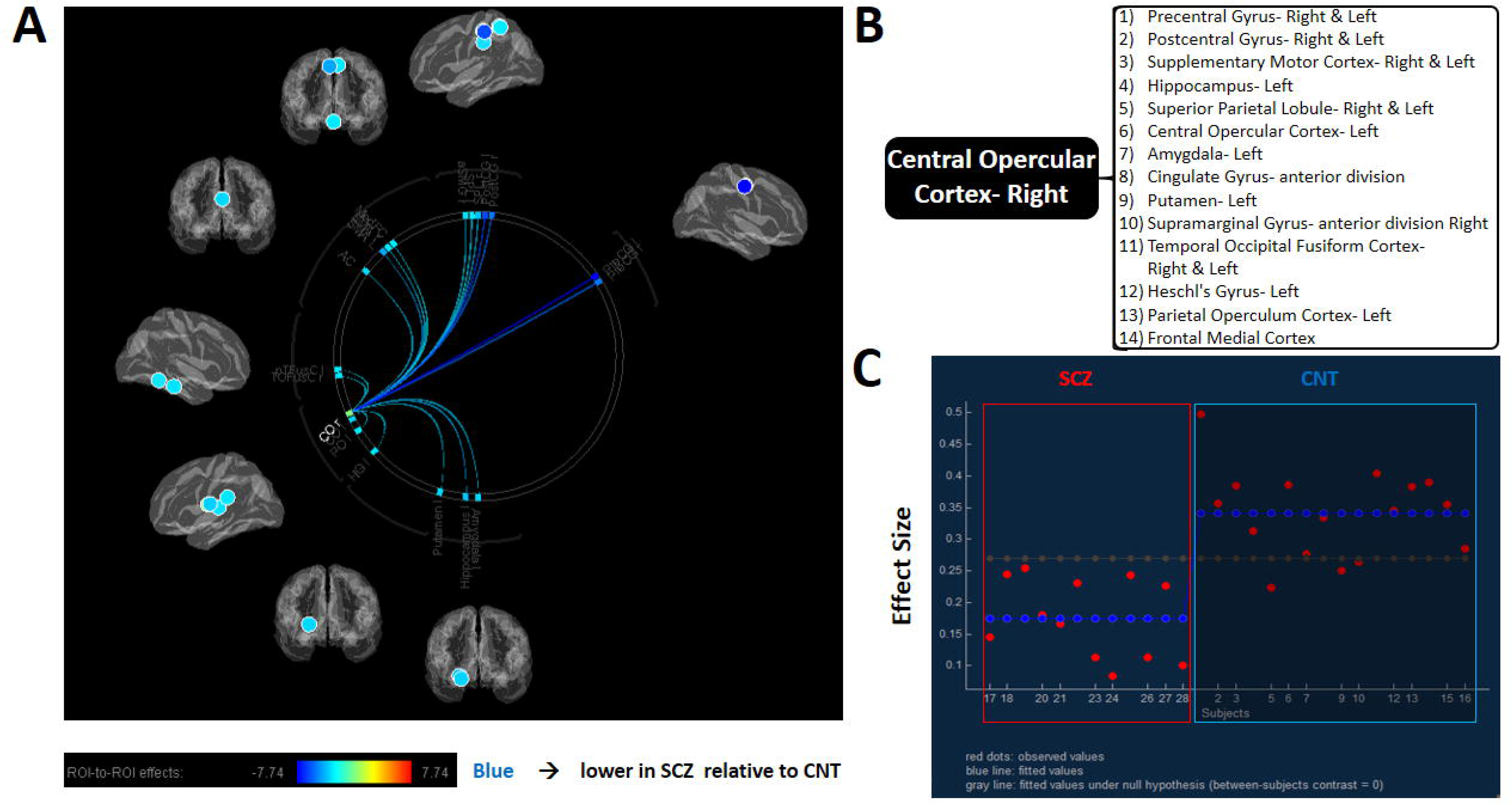
**Functional connectivity difference at region of interest level (ROI-ROI)** between healthy controls (CNT) and patients with schizophrenia (SCZ) during gestalt-perception and corollary discharge tasks combined. (A) Connectivity ring showing the seed ROI and its connected ROIs, that were significantly reduced in patients with schizophrenia (blue indicate reduction and its intensity indicates magnitude). (B) List of connected ROIs. (C) Effect size measures (arbitrary units) across this network for each subject (patients highlighted by red square). Both connection-level thresholds and network-level thresholds applied with FDR corrections.

Both gestalt-perception and corollary-discharge conditions were considered together as a measure of error prediction for ROI-ROI based fMRI connectivity analysis. PSZ showed significantly weaker connectivity for right central opercular cortex with 14 other brain regions (Table-4).

**Table 4:** ROI-ROI connectivity difference between Schizophrenia and Controls during ‘ANGEL’ task in fMRI.

| ROIs connected to Seed ROI<br>(right Central Opercular cortex) | Statistic<br>[F(6,21)=10.35;<br>Intensity=75.31;<br>Size = 19] | Uncorrected<br>p-value | FDR-corrected<br>p-value |
| --- | --- | --- | --- |
| Precentral Gyrus- Left | T(26) = -7.74 | 0 | 0 |
| Postcentral Gyrus – Left | T(26) = -6.19 | 0 | 0.0001 |
| Precentral Gyrus- Right | T(26) = -5.44 | 0 | 0.0005 |
| Precentral Gyrus- Right | T(26) = -5.30 | 0 | 0.0005 |
| Supplementary Motor Cortex – Left | T(26) = -4.33 | 0.0002 | 0.0054 |
| Hippocampus – Left | T(26) = -3.91 | 0.0006 | 0.0122 |
| Superior Parietal Lobule – Right | T(26) = -3.88 | 0.0006 | 0.0122 |
| Central Opercular Cortex – Left | T(26) = -3.55 | 0.0015 | 0.0252 |
| Amygdala – Left | T(26) = -3.44 | 0.002 | 0.0266 |
| Anterior Cingulate | T(26) = -3.42 | 0.0021 | 0.0266 |
| Putamen – Left | T(26) = -3.40 | 0.0022 | 0.0266 |
| Anterior Supramarginal Gyrus –Right | T(26) = -3.29 | 0.0029 | 0.0324 |
| Temporal Occipital Fusiform Cortex – Right | T(26) = -3.14 | 0.0042 | 0.0386 |
| Primary Auditory Cortex – Left | T(26) = -3.12 | 0.0044 | 0.0386 |
| Temporal Occipital Fusiform Cortex – Left | T(26) = -3.11 | 0.0046 | 0.0386 |
| Parietal Operculum Cortex – Left | T(26) = -3.10 | 0.0046 | 0.0386 |
| Supplementary Motor Cortex – Right | T(26) = -3.04 | 0.0054 | 0.0426 |
| Medial Frontal Cortex | T(26) = -2.96 | 0.0064 | 0.0467 |
| Superior Parietal Lobule – Left | T(26) = -2.95 | 0.0066 | 0.0467 |
| **NOTE:** The statistical values were thresholded using a combination of connection-level thresholds (false discovery rate (FDR) estimations with the level of significance set a priori at $p < 0.05$) and network-level thresholds (network-based statistics using permutation tests in conjunction with FDR corrections set at $p < 0.05$), both two-sided. | | | |

#### b) ROI-Voxel analysis (Fig-10)

**Fig-10:**
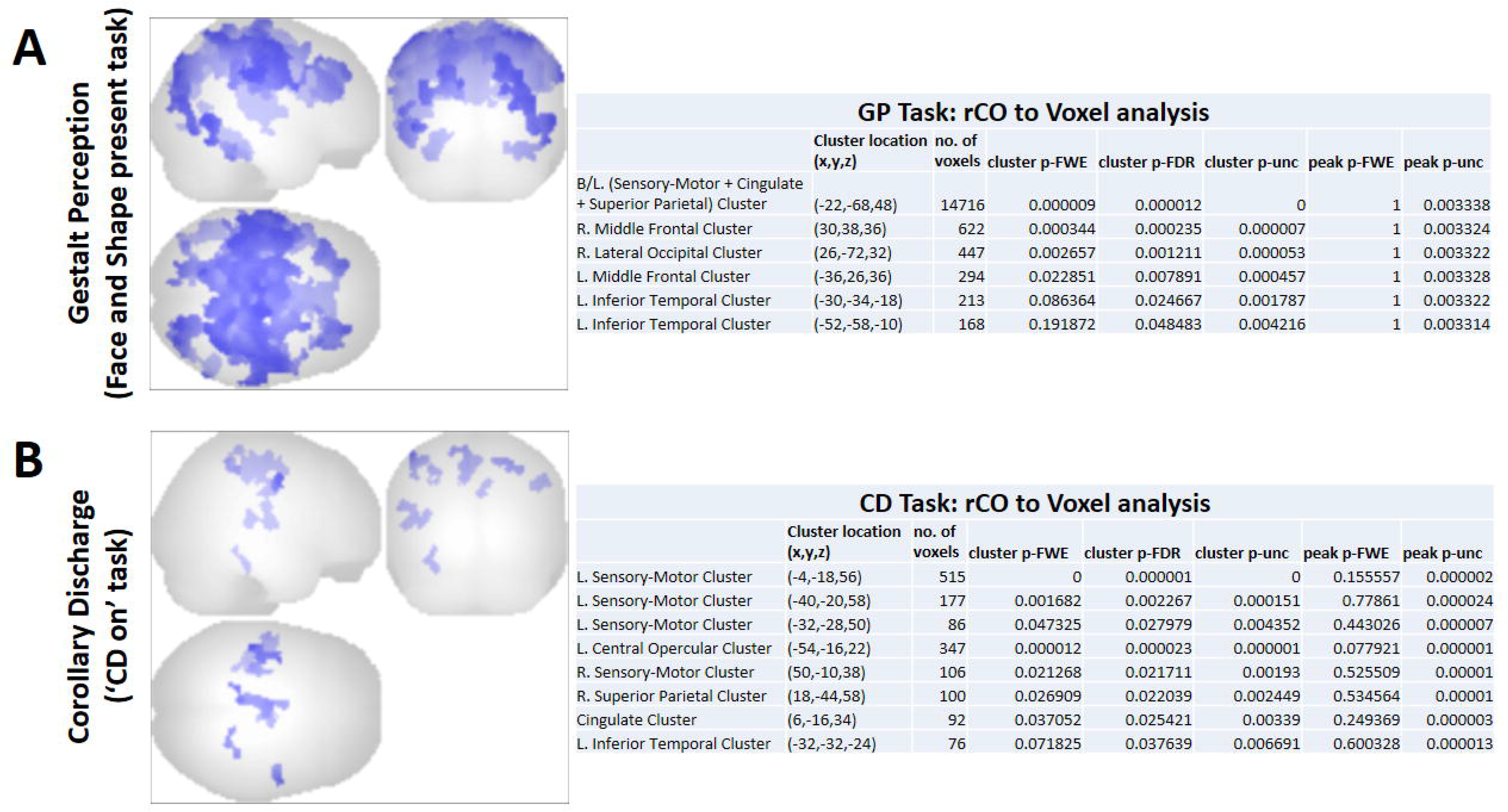
**Functional connectivity difference of right central opercular cortex with all other brain voxels (ROI-Voxel)** between healthy controls (CNT) and patients with schizophrenia (SCZ) during: (A) Gestalt-perception (GP), and (B) Corollary discharge (‘CD on’). Three view glass brains on the right show the voxel clusters that showed significant reduction in connectivity with right central opercular cortex (rCO) during either of tasks, among the patients. The parameters of the identified voxel clusters and level of significance are shown on the left side of each panel. The statistical values were thresholded using a combination of intensity-level thresholds (false discovery rate (FDR) estimations with the level of significance set a priori at p<0.05) and cluster-level thresholds (FDR corrections set at p<0.05), both one-sided (negative).

ROI-Voxel analysis using right central opercular cortex ROI as seed, showed the difference in connected brain regions between task conditions. During both gestalt-perception (face present and shape present trials) and corollary-discharge (‘CDon’ trials), clusters were identified in bilateral sensory-motor, cingulate, right superior parietal and left inferior temporal areas, all showing reduced connectivity in PSZ. Gestalt-perception condition showed many more poorly connected regions than corollary-discharge condition, especially in clusters around bilateral middle frontal areas and right lateral occipital areas. During disrupted gestalt-perception (face absent and shape absent trials) and absent corollary-discharge (‘CDoff’ trials) conditions, no brain areas showed significant difference between the groups.

## Discussion

In the present study, patients with schizophrenia demonstrated various aspects of prediction-errors in terms of altered ERPs and associated measures of EEG dynamics (ERSP and ITC), during both gestalt-perception and corollary-discharge. Such prediction-errors reflect defective brain oscillatory events and thalamo-cortical dysfunctions associated with schizophrenia.

Patients showed reduced N170 amplitude in bilateral parieto-occipital electrodes as well as in frontal electrodes associated with gestalt-perception. In addition to ‘Mooney faces’, N170 changes were also seen for non-face visual-patterns such as the ‘Kanizsa triangle’. This reflects involvement of brain areas outside face-selective network such as medial fusiform gyrus^27^ and superior temporal sulcus^28^. In line with our ERP findings in PSZ, Spencer et al^29^ had found deficits in P1 and N1 components while viewing illusory squares, whereas Turetsky et al^30^ reported N170 deficit for images with facial expressions. N170, occurring 170ms post stimulus-onset in lateral parieto-occipital electrodes, is thought to reflect activation of brain regions processing face-specific visual-patterns (occipital face area, fusiform face area and superior temporal sulcus)^31, 32^. Reduced N170 amplitude in PSZ could be associated with reduced activity in the above-mentioned brain areas. Moreover, it could also result from deficits in long-range brain connectivity in PSZ, as previous EEG studies^29, 33^ have reported reduced stimulus-induced synchronisation in beta and gamma oscillations during gestalt-perception. Interestingly, our PSZ showed deficits in theta frequency, especially in the post stimulus 200ms components of ERSP (over most of scalp electrodes) and ITC deficits observed at peri-stimulus period (mainly over frontal electrodes). Though we did not replicate deficits in higher frequencies, authors of the previous study attributed the deficit in gamma synchronisation to dysfunctional thalamocortical circuitry and hinted on a possible interaction with low frequency oscillations. Furthermore, poor theta coherence among the patients have been implicated in impaired perceptual experience^34^. Theta oscillation over medial frontal scalp sites (presumably generated by medial frontal area) is implicated in cognitive monitoring and thereby contributing to conflict monitoring and sustained attention^35^. It is assumed that medial frontal theta oscillations drives the lateral frontal cortex into synchrony with multiple task-related brain networks^35–37^. This is generally seen as an increase in task-related theta power in frontal brain areas (corresponding to medial to lateral frontal communication) and increase in theta phase synchronization across task-relevant brain networks. Therefore, the deficit in theta oscillation among PSZ would also indicate aberrant neural synchrony, as described using higher oscillations in previous studies.

Patients also showed deficits in ERP morphology at time-windows preceding the stimulus as well as before the N100-P200 ERP complex generated during the corollary-discharge task (‘CDon’ only). These deficits were mainly observed over the centro-parietal electrodes. Patients however did not show suppression of N100-P200 complex as reported in earlier studies^11, 38^. In these studies, suppression of N1 (or N100) component of auditory ERP to sound produced while talking or on button-press was reduced in PSZ. These studies hypothesised that normally, the secondary events of corollary-discharge are associated with sensory attenuation of auditory feedbacks. Accordingly, the conventional ERP components (N100 and P200) would have been chosen in these earlier studies. As we did not observe such a suppression deficit in PSZ, their corollary-discharge deficit could be weaker than reported by prior studies. In the current study, corollary-discharge was studied while participants were performing a more engaging multi-stimuli ANGEL task^22^. The numerous factors that are part of such a task (like multi-sensory integration, motivation, attention, distraction, etc.) could have improved brain response^39^, and allowed prediction-errors to be optimal for sensory attenuation as prediction-errors are also dependent on top-down influences (such as expectations, prior experience, etc.)^3^. In line with the ERP findings, event-related time-frequency analysis in PSZ also found significant deficits at time points before N100-P200 complex for theta oscillations, in terms of magnitude (ERSP) as well as phase (ITC) synchronization. An increase in theta coherence in the pre-stimulus period is known to be associated with corollary-discharge, and found abnormal in PSZ^40^. We also found the topographic distribution of theta dynamics to be severely limited in PSZ, in agreement with earlier reports of poor fronto-temporal theta coherence^40^. Taken together, our ERP, ERSP and ITC findings could indeed reflect corollary-discharge deficits among PSZ, complementing the previous studies.

In our study, PSZ patients showed deficits in many of the event-related EEG dynamics, including highly reduced ERPs and altered theta dynamics, during both gestalt-perception and corollary-discharge. Both gestalt-perception and corollary-discharge reflect the brain’s error prediction mechanisms, and hence altered event-related EEG dynamics seen in the current study could be linked to altered trans-thalamic (thalamo-cortical, including cortico-thalamo-cortical) oscillatory mechanisms^41, 42^. The magnitude of event-related EEG dynamics indicate the integrity of thalamo-cortical network^43^, and theta synchronization is linked to activity of higher order thalamic nuclei^44^. Therefore, our observation of wide spread deficits in scalp distribution of event-related EEG changes, is suggestive of altered connectivity and modulatory changes among remote brain areas. Studies have shown that lesions in dorsolateral prefrontal cortex reduce the visual N100 (or N170) amplitude of extrastriate occipito-temporal areas^45, 46^ and lesions in noradrenergic brain stem modulators cause a reduction in the P3 (or P300) amplitude^47^. Our study thus highlight aberrant neural synchrony during gestalt-perception as well as corollary-discharge associated with schizophrenia supporting the thalamo-cortical dysfunctioning hypothesis^12^.

EEG source-level analysis demonstrated the brain regions involved in such aberrant EEG synchrony associated with schizophrenia. During gestalt-perception task, PSZ showed higher activity in some EEG source clusters (ERP in right anterior cingulate; with increased theta-ERSP and decreased beta-ERSP in left superior parietal; theta-ITC in left anterior cingulate), and lower activity in other clusters (ERP in left motor; theta-ERSP increase and beta-ERSP decrease in right insular; theta-ITC in left frontal opercular). Though there are no known reports on EEG source-level studies of gestalt-perception deficits in schizophrenia, magneto-encephalographic (MEG) studies in non-clinical^48^ and autism-spectrum-disorder^49^ population suggest that impaired gestalt-perception is associated with aberrant activity in the right fronto-parietal network^50^. Similarly, decreased activity in the dorsolateral prefrontal, medial frontal and parietal cortices have been shown to be associated with gestalt-perception of sematic context in schizophrenia^51^. Insular network which forms part of the larger fronto-parietal network, is known to be active during perception at an abstract level^52–, 54^ and hence could play a core role in gestalt-perception. In addition to the already described role of theta coherence^34^, post-stimulus beta power decrease^9, 29^ is also considered important for gestalt-perception. Taken together, the EEG source analysis highlighting increased theta power and decreased beta power in right insular cluster could indicate gestalt-perception, which is impaired in schizophrenia. The increased theta phase synchrony in left anterior cingulate cluster and theta power increase in left superior parietal cluster among patients could be a futile compensatory drive to activate insular region^35^.

During corollary-discharge task, patients showed deficits in most of the EEG source clusters identified (ERP in left superior temporal; theta-ERSP in left basal ganglia and right superior parietal; theta-ITC in left motor and right cingulate). Moreover, patients showed increased beta power (ERSP) in the right superior parietal cluster suggesting impaired suppression of sensory processing^55, 56^. These results suggest that corollary-discharge involves synchronous activation of frontal (motor) and parietal (somatosensory) regions with mediation from basal ganglia and the anterior cingulate. Impairment in this synchrony would have prevented suppression of sensory processing areas. These are also in agreement with prior reports of a deficit in efference copy during corollary-discharge^38, 40^, where a decreased fronto-temporal theta coherence was observed. As described earlier, theta oscillation deficit could be a cause for this disharmony in patients^35^.

Our fMRI findings highlight the prominent role of cingulo-opercular (or cingulo-insular) network in both gestalt-perception and corollary-discharge, which is significantly affected in PSZ. Component brain regions of this network, especially the insula and anterior cingulate cortex, are believed to play significant roles in error processing, brain state switching, sensory integration and self-awareness^57–59^. As our cognitive task predominantly consists of these components, it is not surprising that cingulo-opercular network connectivity dominates during the task. Moreover, the significant reduction in insular connectivity among PSZ is a widely reported finding, especially in recent studies^60, 61^. In the light of our recent finding of sleep deficits in Schizophrenia^62^, we would have expected a larger impact on thalamo-cortical connectivity. Interestingly, a recent fMRI study reported thalamic hyper-connectivity and sensory hypo-connectivity in schizophrenia during resting state^63^. They relate this to the differential connectivity of thalamus seen among patients^64–66^, with stronger connectivity between thalamus and sensory-motor cortices and weaker connectivity between thalamus and prefrontal cortex. It is speculated that such an aberrant connectivity during resting state would correspond to cortical-subcortical antagonism resulting in break-down of default-mode network or even descent into light sleep. Thus, delinking of insula (or its expanded cingulo-opercular network) and higher-order thalamic nuclei^67^ could explain the prediction-error and poor self-monitoring that is a hallmark of schizophrenia.

### Limitations

As our EEG source localization relied on default channel locations (instead of individual digitized locations) and template MRIs (instead of individual MRI images), there could be localization errors in the EEG-sources we reported. However, our EEG and fMRI findings show common dysfunctional brain areas in PSZ and the brain regions identified conforms to those reported in literature; thus, our results seem robust to the above limitation. We could not do the EEG and fMRI study on the same group, but in turn, this gave us an opportunity to replicate the findings in an independent sample.

## Conclusion

This is the first study to examine prediction-error-coding during both perception (gestalt-perception) and action (corollary-discharge) within the purview of ‘thalamo-cortical dysfunction hypothesis’ in schizophrenia. We identified deficits in ERPs, theta-oscillations and localised abnormality in insula-driven brain network, using separate high-density EEG and fMRI studies. Besides adding to the knowledge-base of schizophrenia research, our novel task design and findings on theta-oscillation could benefit in the development of effective neuromodulatory therapeutic tools for PSZ such as neurofeedback and transcranial brain stimulation.

## Supporting information

supplementary methodology

## Acknowledgements

This work was funded in part by Indian Council for Medical Research (ICMR), Government of India (Senior Research Fellowship; Ref. No.: 3/1/3/37/Neuro/2013-NCD-I to A.S.) and Department of Biotechnology (DBT), Government of India (Grant No. BT/PR/8363/MED/14/1252 to J.P.J).

## References

1. Insel T, Cuthbert B, Garvey M, et al. Research domain criteria (RDoC): toward a new classification framework for research on mental disorders. Am J Psychiatry. 2010;167:748–51.

2. Friston K. Prediction, perception and agency. Int J Psychophysiol. 2012;83(2):248-252. doi:10.1016/j.ijpsycho.2011.11.014.

3. Lawson RP, Rees G, Friston KJ. An aberrant precision account of autism. Front Hum Neurosci. 2014;8(May):302. doi:10.3389/fnhum.2014.00302.

4. Koster-Hale J, Saxe R. Theory of Mind: A Neural Prediction Problem. Neuron. 2013;79(5):836–848. doi:10.1016/j.neuron.2013.08.020.

5. Feinberg I. Efference copy and corollary discharge: implications for thinking and its disorders. Schizophr Bull. 1978;4(4):636–640. doi:10.1093/schbul/4.4.636.

6. Ford JM, Mathalon DH. Anticipating the future: Automatic prediction failures in schizophrenia. Int J Psychophysiol. 2012;83(2):232–239. doi:10.1016/j.ijpsycho.2011.09.004.

7. Yamashita Y, Tani J. Spontaneous prediction error generation in schizophrenia. PLoS One. 2012;7(5):e37843. doi:10.1371/journal.pone.0037843.

8. Uhlhaas PJ, Mishara AL. Perceptual anomalies in schizophrenia: Integrating phenomenology and cognitive neuroscience. Schizophr Bull. 2007;33(1):142–156. doi:10.1093/schbul/sbl047.

9. Spencer KM, Ghorashi S. Oscillatory dynamics of Gestalt perception in schizophrenia revisited. Front Psychol. 2014;5(February):68. doi:10.3389/fpsyg.2014.00068.

10. Ford JM, Gray M, Faustman WO, Roach BJ, Mathalon DH. Dissecting corollary discharge dysfunction in schizophrenia. Psychophysiology. 2007;44(4):522–529. doi:10.1111/j.1469-8986.2007.00533.x.

11. Ford JM, Palzes V A., Roach BJ, Mathalon DH. Did i do that? Abnormal predictive processes in schizophrenia when button pressing to deliver a tone. Schizophr Bull. 2014;40(4):804–812. doi:10.1093/schbul/sbt072.

12. Vukadinovic Z, Rosenzweig I. Abnormalities in thalamic neurophysiology in schizophrenia: Could psychosis be a result of potassium channel dysfunction? Neurosci Biobehav Rev. 2012;36(2):960–968. doi:10.1016/j.neubiorev.2011.11.005.

13. Lisman JE. Excitation, inhibition, local oscillations, or large-scale loops: What causes the symptoms of schizophrenia? Curr Opin Neurobiol. 2012;22(3):537–544. doi:10.1016/j.conb.2011.10.018.

14. Vukadinovic Z. Sleep abnormalities in schizophrenia may suggest impaired trans-thalamic cortico-cortical communication: Towards a dynamic model of the illness. Eur J Neurosci. 2011;34(7):1031–1039. doi:10.1111/j.1460-9568.2011.07822.x.

15. Ferrarelli F, Tononi G. The thalamic reticular nucleus and schizophrenia. Schizophr Bull. 2011;37(2):306–315. doi:10.1093/schbul/sbq142.

16. Ferrarelli F, Peterson MJ, Sarasso S, et al. Thalamic dysfunction in schizophrenia suggested by whole-night deficits in slow and fast spindles. Am J Psychiatry. 2010;167(11):1339–1348. doi:10.1176/appi.ajp.2010.09121731.

17. Wamsley EJ, Tucker M A., Shinn AK, et al. Reduced sleep spindles and spindle coherence in schizophrenia: Mechanisms of impaired memory consolidation? Biol Psychiatry. 2012;71(2):154–161. doi:10.1016/j.biopsych.2011.08.008.

18. Onitsuka T, Oribe N, Nakamura I, Kanba S. Review of neurophysiological findings in patients with schizophrenia. Psychiatry Clin Neurosci. 2013;67(7):461–470. doi:10.1111/pcn.12090.

19. Williams TJ, Nuechterlein KH, Subotnik KL, Yee CM. Distinct neural generators of sensory gating in schizophrenia. Psychophysiology. 2011;48(4):470–478. doi:10.1111/j.1469-8986.2010.01119.x.

20. Buchmann A, Dentico D, Peterson MJ, et al. Reduced mediodorsal thalamic volume and prefrontal cortical spindle activity in schizophrenia. Neuroimage. 2014;102(2):540–547. doi:10.1016/j.neuroimage.2014.08.017.

21. Woodward ND, Karbasforoushan H, Heckers S. Thalamocortical dysconnectivity in schizophrenia. Am J Psychiatry. 2012;169(October):1092–1099. doi:10.1176/appi.ajp.2012.12010056.

22. Nair AK, Sasidharan A, John JP, Mehrotra S, Kutty BM. Assessing neurocognition via gamified experimental logic: A novel approach to simultaneous acquisition of multiple ERPs. Front Neurosci. 2016;10(January):1–14. doi:10.3389/fnins.2016.00001.

23. Delorme A, Makeig S. EEGLAB: An open source toolbox for analysis of single-trial EEG dynamics including independent component analysis. J Neurosci Methods. 2004;134:9–21. doi:10.1016/j.jneumeth.2003.10.009.

24. Mullen T, Kothe C, Chi YM, et al. Real-time modeling and 3D visualization of source dynamics and connectivity using wearable EEG. In: Conference Proceedings: Annual International Conference of the IEEE Engineering in Medicine and Biology Society. IEEE Engineering in Medicine and Biology Society. Conference. Vol 2013. NIH Public Access; 2013:2184-2187. doi:10.1016/j.biotechadv.2011.08.021.Secreted.

25. Bigdely-Shamlo N, Mullen T, Kreutz-Delgado K, Makeig S. Measure projection analysis: A probabilistic approach to EEG source comparison and multi-subject inference. Neuroimage. 2013;72:287–303. doi:10.1016/j.neuroimage.2013.01.040.

26. Whitfield-Gabrieli S, Nieto-Castanon A. Conn: A Functional Connectivity Toolbox for Correlated and Anticorrelated Brain Networks. Brain Connect. 2012;2(3):125–141. doi:10.1089/brain.2012.0073.

27. Sadeh B, Yovel G. Why is the N170 enhanced for inverted faces? An ERP competition experiment. Neuroimage. 2010;53(2):782–789. doi:10.1016/j.neuroimage.2010.06.029.

28. Itier RJ, Taylor MJ. Source analysis of the N170 to faces and objects. Neuroreport. 2004;15(8):1261–1265. doi:10.1097/01.wnr.0000127827.73576.d8.

29. Spencer KM, Nestor PG, Perlmutter R, et al. Neural synchrony indexes disordered perception and cognition in schizophrenia. Proc Natl Acad Sci U S A. 2004;101(49):17288–17293. doi:10.1073/pnas.0406074101.

30. Turetsky BI, Kohler CG, Indersmitten T, Bhati MT, Charbonnier D, Gur RC. Facial emotion recognition in schizophrenia:When and why does it go awry? Schizophr Res. 2007;94(1-3):253–263. doi:10.1016/j.schres.2007.05.001.

31. Rossion B, Gauthier I. How does the brain process upright and inverted faces? Behav Cogn Neurosci Rev. 2002;1(1):63–75. doi:10.1177/1534582302001001004.

32. Nguyen VT, Cunnington R. The superior temporal sulcus and the N170 during face processing: Single trial analysis of concurrent EEG-fMRI. Neuroimage. 2014;86:492–502. doi:10.1016/j.neuroimage.2013.10.047.

33. Uhlhaas PJ, Linden DEJ, Singer W, et al. Dysfunctional long-range coordination of neural activity during Gestalt perception in schizophrenia. J Neurosci. 2006;26(31):8168–8175. doi:10.1523/JNEUROSCI.2002-06.2006.

34. Slagter HA, Lutz A, Greischar LL, Nieuwenhuis S, Davidson RJ. Theta Phase Synchrony and Conscious Target Perception: Impact of Intensive Mental Training. J Cogn Neurosci. 2009;21(8):1536–1549. doi:10.1162/jocn.2009.21125.Theta.

35. Clayton MS, Yeung N, Cohen Kadosh R. The roles of cortical oscillations in sustained attention. Trends Cogn Sci. 2015;19(4):188–195. doi:10.1016/j.tics.2015.02.004.

36. Cohen MX. A neural microcircuit for cognitive conflict detection and signaling. Trends Neurosci. 2014;37(9):480–490. doi:10.1016/j.tins.2014.06.004.

37. Oehrn CR, Hanslmayr S, Fell J, et al. Neural Communication Patterns Underlying Conflict Detection, Resolution, and Adaptation. J Neurosci. 2014;34(31):10438–10452. doi:10.1523/JNEUROSCI.3099-13.2014.

38. Mathalon DH, Ford JM. Corollary discharge dysfunction in schizophrenia: evidence for an elemental deficit. Clin EEG Neurosci. 2008;39(2):82–86. doi:10.1177/155005940803900212.

39. Wynn JK, Jahshan C, Green MF. Multisensory integration in schizophrenialJ: a behavioural and event-related potential study. Cogn Neuropsychiatry. 2015;19(4):319–336. doi:10.1080/13546805.2013.866892.

40. Ford JM, Mathalon DH, Whitfield S, Faustman WO, Roth WT. Reduced communication between frontal and temporal lobes during talking in schizophrenia. Biol Psychiatry. 2002;51(6):485–492. doi:10.1016/S0006-3223(01)01335-X.

41. Vukadinovic Z. NMDA receptor hypofunction and the thalamus in schizophrenia. Physiol Behav. 2014;131:156–159. doi:10.1016/j.physbeh.2014.04.038.

42. Sherman SM, Guillery RW. Distinct functions for direct and transthalamic corticocortical connections. J Neurophysiol. 2011;106(June 2011):1068–1077. doi:10.1152/jn.00429.2011.

43. Näätänen R, Picton T. The N1 wave of the human electric and magnetic response to sound: a review and an analysis of the component structure. Psychophysiology. 1987;24:375–425. doi:10.1111/j.1469-8986.1987.tb00311.x.

44. Saalmann YB, Pinsk MA, Wang L, Li X, Kastner S. The Pulvinar Regulates Information Transmission Between Cortical Areas Based on Attention Demands. Science (80-). 2012;337(6095):753-756. doi:10.1126/science.1223082.

45. Barceló F, Suwazono S, Knight RT. Prefrontal modulation of visual processing in humans. Nat Neurosci. 2000;3(4):399–403. doi:10.1038/73975.

46. Swick D, Knight RT. Cortical lesions and attention. In: Parasuraman R, ed. The Attentive Brain. Cambridge, MA: MIT Press; 1998:143-162.

47. Pineda JA, Foote SL, Neville HJ. Effects of locus coeruleus lesions on auditory, long-latency, event-related potentials in monkey. J Neurosci. 1989;9(1):81–93.

48. Grützner C, Uhlhaas PJ, Genc E, Kohler A, Singer W, Wibral M. Neuroelectromagnetic correlates of perceptual closure processes. J Neurosci. 2010;30(24):8342–8352. doi:10.1523/JNEUROSCI.5434-09.2010.

49. Sun L, Grutzner C, Bolte S, et al. Impaired Gamma-Band Activity during Perceptual Organization in Adults with Autism Spectrum Disorders: Evidence for Dysfunctional Network Activity in Frontal-Posterior Cortices. J Neurosci. 2012;32(28):9563–9573. doi:10.1523/JNEUROSCI.1073-12.2012.

50. Uhlhaas PJ, Singer W. Neuronal Dynamics and Neuropsychiatric Disorders: Toward a Translational Paradigm for Dysfunctional Large-Scale Networks. Neuron. 2012;75(6):963–980. doi:10.1016/j.neuron.2012.09.004.

51. Kuperberg GR, West WC, Lakshmanan BM, Goff D. Functional magnetic resonance imaging reveals neuroanatomical dissociations during semantic integration in schizophrenia. Biol Psychiatry. 2008;64(5):407–418. doi:S0006-3223(08)00362-4 [pii] 10.1016/j.biopsych.2008.03.018.

52. Tamura Y, Kuriki S, Nakano T. Involvement of the left insula in the ecological validity of the human voice. Sci Rep. 2015;5:8799. doi:10.1038/srep08799.

53. Craig ADB. How do you feel--now? The anterior insula and human awareness. Nat Rev Neurosci. 2009;10(1):59–70. doi:10.1038/nrn2555.

54. Wong PCM, Parsons LM, Martinez M, Diehl RL. The role of the insular cortex in pitch pattern perception: the effect of linguistic contexts. J Neurosci. 2004;24(41):9153–9160. doi:10.1523/JNEUROSCI.2225-04.2004.

55. Engel AK, Fries P. Beta-band oscillations-signalling the status quo? Curr Opin Neurobiol. 2010;20(2):156–165. doi:10.1016/j.conb.2010.02.015.

56. Waldhauser GT, Johansson M, Hanslmayr S. Alpha/Beta Oscillations Indicate Inhibition of Interfering Visual Memories. J Neurosci. 2012;32(6):1953–1961. doi:10.1523/JNEUROSCI.4201-11.2012.

57. Tang Y-YY, Rothbart MK, Posner MI. Neural correlates of establishing, maintaining, and switching brain states. Trends Cogn Sci. 2012;16(6):997–1003. doi:10.1016/j.tics.2012.05.001.

58. Seth AK, Suzuki K, Critchley HD. An interoceptive predictive coding model of conscious presence. Front Psychol. 2012;3(January):395. doi:10.3389/fpsyg.2011.00395.

59. Klein T A, Ullsperger M, Danielmeier C. Error awareness and the insula: links to neurological and psychiatric diseases. Front Hum Neurosci. 2013;7(February):14. doi:10.3389/fnhum.2013.00014.

60. Moran L V., Tagamets MA, Sampath H, et al. Disruption of anterior insula modulation of large-scale brain networks in schizophrenia. Biol Psychiatry. 2013;74(6):467–474. doi:10.1016/j.biopsych.2013.02.029.

61. Palaniyappan L, Simmonite M, White TP, Liddle EB, Liddle PF. Neural primacy of the salience processing system in schizophrenia. Neuron. 2013;79(4):814–828. doi:10.1016/j.neuron.2013.06.027.

62. Sasidharan A, Kumar S, Nair AK, et al. Further evidences for sleep instability and impaired spindle-delta dynamics in schizophrenia: a whole-night polysomnography study with neuroloop-gain and sleep-cycle analysis. Sleep Med. 2017. doi:10.1016/j.sleep.2017.02.009.

63. Damaraju E, Allen EA, Belger A, et al. Dynamic functional connectivity analysis reveals transient states of dysconnectivity in schizophrenia. NeuroImage Clin. 2014;5(July):298–308. doi:10.1016/j.nicl.2014.07.003.

64. Woodward ND, Karbasforoushan H, Heckers S. Thalamocortical dysconnectivity in schizophrenia. Am J Psychiatry. 2012;169(10):1092–1099. doi:10.1176/appi.ajp.2012.12010056.

65. Anticevic A, Cole MW, Repovs G, et al. Characterizing Thalamo-Cortical Disturbances in Schizophrenia and Bipolar Illness. Cereb Cortex. 2014. doi:10.1093/cercor/bht165.

66. Marenco S, Stein JL, Savostyanova AA, et al. Investigation of Anatomical Thalamo-Cortical Connectivity and fMRI Activation in Schizophrenia. Neuropsychopharmacology. 2012;37:499–507. doi:10.1038/npp.2011.215.

67. Corradi-Dell’Acqua C, Tomelleri L, Bellani M, et al. Thalamic-insular dysconnectivity in schizophrenia: Evidence from structural equation modeling. Hum Brain Mapp. 2012;33(3):740–752. doi:10.1002/hbm.21246.

