## supplementary methodology for "Schizophrenia patients show aberrant brain dynamics associated with gestalt-perception and corollary discharge: Reflections from ERP and fMRI findings"

Supplementary Methods

### Participants

### Only right-handed participants (as determined by Edinburgh’s inventory) were included in the study. The diagnosis of schizophrenia was arrived at using criteria from DSM–IV (1) based on the consensus of a research psychiatrist who conducted a semi-structured interview and a trained research psychologist who used the Mini International Neuropsychiatric Interview for DSM-IV (MINI-Plus) (2). In case of individuals in the control group, presence of any medical/psychiatric/neurological condition requiring continuous medications, current psychotropic use and history of psychiatric illness in first-degree relatives were ruled out by an unstructured clinical interview.

### Task design

### In brief, the subject watches a pair of two-tone images (familiar pattern and checker pattern) on a screen and must press a left or right button based on simple rules for the familiar pattern (changes with game-level). There are three game-levels with increasing complexity, each game-level is about 15 minutes and overall time for EEG-ERP acquisition is about 1.5 hours. The familiar pattern could be from a set of degraded human faces (Mooney face; ‘Face present’ trials) or illusory shapes (Kanizsa triangle; ‘Shape present’ trials), both testing the grouping of basic image elements into perception of a global gestalt (3). Scrambled counterparts of these images (‘Face absent’ and ‘Shape absent’ trials) were also used to evoke more local perception in contrast to gestalt perception. Corollary discharge mechanism requires a motor response and its sensory feedback to occur with a time interval as predicted from the motor act, so as to tag it as ‘self-generated’ (4). This was implemented during the game levels 2 and 3, where an auditory feedback (60ms long polysyllabic tone) is provided with each button response (corollary discharge present or ‘CDon’ trials). At times a random delay is introduced between the feedback and button press, and this incongruity disrupts corollary discharge mechanism (corollary discharge absent or ‘CDoff’ trials) to tag it as ‘other-generated’ trials. The trials of game level 1 having button presses without feedback would act as control condition for the corollary discharge trials.

The shortened version used for fMRI study had a single level with components of game-levels 1 and 2 combined, less number of trials per block (20 instead of 25), longer inter-trial interval to match scanning interval (2s instead of 1.5s), and thus having a duration of about 14 minutes.

### EEG recording

### The study was carried out either in the afternoons (2-6pm) or mornings (8-12am), counterbalanced between the subjects. Ambient temperature maintained at 25^o^C. 62 monopolar electrodes for EEG, and 2 sets of bipolar electrodes for electro-oculogram (EOG) were used. EEG electrode positions were in accordance with the 10-10 international system for electrode placement. Reference electrode was between Cz and CPz, and ground between Fz and FPz. Horizontal EOG electrodes were placed 2cm away from the outer canthi of both the eyes, and vertical EOG, electrodes were placed 2cm above and below the left eye. Disposable syringes (5ml) with 14G blunt needles were used for applying gel. Impedance for all electrodes was maintained between 5 to 10KΩ.

### ERP analysis

EEG drift artefacts removed using a 0.25-0.75Hz transition band high-pass filter, bad channels were rejected using an automated correlation based routine and high amplitude stereotypical (e.g., eye blinks) and non-stereotypical (e.g., movement) artefacts removed using an artefact subspace reconstruction (ASR) algorithm (5); all implemented as a plugin in EEGLAB. ASR relies on a sliding window principal component analysis (PCA) to statistically compare with a reference clean portion of the EEG data and then linearly reconstruct deviant portions of the data (‘artefact subspaces’) based on the correlation structure observed in the reference clean data. We used a threshold of 5 standard deviations for ASR.

ERP waveform time-locked to gestalt-perception images (‘Face absent’, ‘Face present’, ‘Shape absent’ and ‘Shape present’) showing a prominent negative (N170) and positive component (P200) at the parieto-occipital scalp site (PO8) formed the validation of the Gestalt-perception task. The N1 and P2 ERP waveforms that are known to be produced during CDoff’ condition and attenuated during ‘CDon’ condition formed the validation of the Corollary-discharge task.

A total of 4094 ‘Face absent’ trials (95.19 ± 17.90 per SCZ; 99.76 ± 1.67 per CNT), 4215 ‘Face present’ trials (101.05 ± 14.30 per SCZ; 99.67 ± 5.19 per CNT), 4000 ‘Shape absent’ trials (92.10 ± 23.03 per SCZ; 98.38 ± 5.50 per CNT), and 4263 ‘Shape present’ trials (102.52 ± 13.21 per SCZ; 100.48 ± 4.74 per CNT) were available for analysis of gestalt perception. A total of 8063 ‘CDon’ trials (190.29 ± 25.39 per SCZ; 193.67 ± 8.18 per CNT) and 8236 ‘CDoff’ trials (194 ± 29.13 per SCZ; 198.19 ± 5.58 per CNT) were available for analysis of corollary discharge mechanism.

### Event-Related Spectral Perturbations (ERSP) and Inter-Trial Coherence (ITC) analysis

ERSP represents the power modulations relative to pre-stimulus period and consistent across trials. Whereas ITC is a measure of strength of phase alignment across trials, relative to stimulus onset. ERSP and ITC involves a trial-by-trial time-frequency analysis of the event-related epochs based on sinusoidal moving Hanning-windowed wavelet with linearly increased cycles (6), from 3 cycles for lowest frequency (3Hz) to 15 cycles for highest frequency (30Hz) analysed. The extracted magnitude and phase information for each frequency across each window is then divided by the root-mean-square value of the corresponding baseline window (the one preceding the stimulus). The ERSP values are scaled in decibels (log transformed) while ITC scale from 0 (weakest) to 1 (strongest).

### EEG Source level analysis

To further examine the brain sources of ERP signals (source localization), first an independent component analysis (ICA) was performed onto the epoched data (-500ms to 800ms relative to image onset; non-overlapping epochs), using the extended ‘runica’ algorithm implemented in EEGLABv13. ICA assumes a linear mixing of brain source signals in the scalp EEG due to volume conduction, and finds a set of spatial filters to unmix the scalp channel data into maximally independent individual time courses having their individual scalp projection patterns (potential information sources). The resulting ICA weight matrix was applied to a longer epoched data (-1000ms to 2000ms relative to gestalt images or corollary discharge sounds) to derive the independent components (ICs) corresponding to each condition (7). Then the location of each IC was computed within a common brain template in the form of equivalent current dipoles using a standard template boundary element model (BEM) in the DIPFIT plugin of EEGLAB (8). Only a subset of ICs with low estimation error (less than 17% residual variance) and those localised within the template brain volume, were used for further analysis. The residual variance cut-off of 17% was decided based on the recommended threshold of 15% by the software and further adjusting this threshold to include at least one IC from each subject for analysis. Measure Projection Analysis (MPA) implemented in MPT (Measure Projection Toolbox) plugin of EEGLAB was used. MPA basically projects the scalp-level IC measures (ERP/ERSP/ITC) to Gaussian-smoothed brain space (using 8mm voxel kernel) estimated from their corresponding dipole locations, and further statistical comparisons are made in this voxel space (9). Next, those projected values associated with highly significant brain locations are then clustered based on their similarities using affinity-propagation method, and these clusters are used for further group-level analysis. The brain locations of these clusters were mapped by the software using LONI Probabilistic Brain Atlas (LPBA40) (10). Default parameters of the MPT plugin were used for the present study; i.e. maximum domain exemplar correlation set at 0.8, and 2000 bootstrap permutations to compute measure convergence with a threshold of p=0.01 (uncorrected).

### fMRI acquisition

The BOLD signals were obtained using a gradient echo-planar imaging (EPI) sequence at a resolution of 3x3x4mm (TR/TE: 2000ms/30ms; flip angle: 78°; slice gap: 0mm; 37 sequential slices in descending order; FOV: 192mm x 192mm; matrix size: 64x64; Parallel imaging: GRAPPA with Acceleration factor 2). Head movements were minimized by using soft memory foam cushions within the head coil. T1-weighted structural images with resolution of 1x1x1mm were also acquired in the same imaging session using a MPRAGE (Magnetization Prepared Rapid Acquisition Gradient Echo) sequence (TR/TE: 1900ms/2.43ms, flip angle: 9°, slice gap: 0mm, 192 interleaved slices, FOV: 256mm x 256mm, matrix size: 256x256, and Parallel imaging: GRAPPA with Acceleration factor 2).

### fMRI Analysis

Before subjecting to the automated analysis routines, the structural and functional images were manually inspected for gross abnormalities/artefacts (e.g., poor image quality, incomplete images, etc.). Additionally, the images were manually aligned to anterior commissure (AC) – posterior commissure (PC) plane with AC as the origin, using SPM software interface (this step was required as SPM analysis routines are sensitive to AC origin discrepancy). Briefly, the automated procedure included the following steps: **(1) Preprocessing**: Rough affine co-registration, Segmentation, Slice time correction, Realignment and Unwarping, Co-registration, Normalization (to the standard Montreal Neurological Institute (MNI) space using the ICBM152 brain template(11, 12); resliced and resampled to a voxel size of 2mm x 2mm x 2mm), Smoothing (8mm full width half maximum or FWHM); **(2) Denoising**: motion parameter regression, removing BOLD CSF and white matter noise, regression of task-induced co-activations (no filters). **(3) A general linear model (GLM) approach**: was adopted to perform subject level correlation analysis between fMRI data from 136 standard regions of interest (ROIs; 95 cortical and 51 subcortical) that are part of CONNv14p Toolbox. The GLM for the connectivity estimations (bivariate correlations) during error prediction trials (both gestalt perception and corollary discharge tasks combined) was queried using appropriate t-contrasts, in comparison to non-task trials. This will cause ROIs in each subject that are more strongly connected during error-prediction task to show up. These are generated in the form of contrast images for individual subjects from each group, and then entered a second level random effects analysis (RFX) for group comparisons. The statistical values of the ROI-ROI connectivity analysis were further thresholded using a combination of connection-level thresholds (false discovery rate (FDR) estimations with the level of significance set a priori at p<0.05) and network-level thresholds (network-based statistics using permutation tests in conjunction with FDR corrections set at p<0.05).

Thus, a whole brain ROI-ROI analysis was used to track down affected brain network without restricting analysis to ROIs set a priori, also reducing the number of pairwise correlation analysis required for such whole-brain assessment. The seed ROI that showed statistical significance in ROI-ROI analysis, was further used in ROI-Voxel analysis (ROI as seed and compared with all other brain voxels during gestalt-perception and corollary-discharge tasks separately), so that the brain areas significantly connected to seed ROI are identified at voxel-level, not restricting to default ROI boundaries.

### References:

1. American Psychiatric Association (2000): *Diagnostic and statistical manual of mental disorders DSM-IV-TR fourth edition (text revision)*. . Washington, DC: American Psychiatric Publishing, Inc.

2. Sheehan D V., Lecrubier Y, Sheehan KH, Amorim P, Janavs J, Weiller E, *et al.* (1998): The Mini-International Neuropsychiatric Interview (M.I.N.I.): The development and validation of a structured diagnostic psychiatric interview for DSM-IV and ICD-10. *J Clin Psychiatry* 59: 22–33.

3. Uhlhaas PJ, Linden DEJ, Singer W, Haenschel C, Lindner M, Maurer K, Rodriguez E (2006): Dysfunctional long-range coordination of neural activity during Gestalt perception in schizophrenia. *J Neurosci* 26: 8168–8175.

4. Ford JM, Palzes V a., Roach BJ, Mathalon DH (2014): Did i do that? Abnormal predictive processes in schizophrenia when button pressing to deliver a tone. *Schizophr Bull* 40: 804–812.

5. Mullen T, Kothe C, Chi YM, Ojeda A, Kerth T, Makeig S, *et al.* (2013): Real-time modeling and 3D visualization of source dynamics and connectivity using wearable EEG. *Conf Proc Annu Int Conf IEEE Eng Med Biol Soc IEEE Eng Med Biol Soc Conf* (Vol. 2013) NIH Public Access, pp 2184–2187.

6. Delorme A, Makeig S (2004): EEGLAB: An open source toolbox for analysis of single-trial EEG dynamics including independent component analysis. *J Neurosci Methods* 134: 9–21.

7. Delorme A, Makeig S (n.d.).: EEGLAB Online tutorial: Decomposing Data Using ICA. Retrieved March 28, 2015, from http://sccn.ucsd.edu/wiki/Chapter_09:_Decomposing_Data_Using_ICA.

8. Oostenveld R, Oostendorp TF (2002): Validating the boundary element method for forward and inverse EEG computations in the presence of a hole in the skull. *Hum Brain Mapp* 17: 179–92.

9. Bigdely-Shamlo N, Mullen T, Kreutz-Delgado K, Makeig S (2013): Measure projection analysis: A probabilistic approach to EEG source comparison and multi-subject inference. *Neuroimage* 72: Elsevier Inc.287–303.

10. Shattuck DW, Mirza M, Adisetiyo V, Hojatkashani C, Salamon G, Narr KL, *et al.* (2008): Construction of a 3D probabilistic atlas of human cortical structures. *Neuroimage* 39: 1064–1080.

11. Mazziotta JC, Toga AW, Evans A, Fox P, Lancaster J (1995): A probabalisitc atlas of the human brain: theory and rationale for its development. *Neuroimage*.

12. Mazziotta J, Toga a, Evans a, Fox P, Lancaster J, Zilles K, *et al.* (2001): A probabilistic atlas and reference system for the human brain: International Consortium for Brain Mapping (ICBM). *Philos Trans R Soc Lond B Biol Sci* 356: 1293–1322.
